# LegoBody: facile generation of bispecific and multi-specific antibodies

**DOI:** 10.1101/2019.12.25.888586

**Authors:** Shanshan Lang, Su Yang, Scott Bidlingmaier, Nam-Kyung Lee, Bin Liu

**Affiliations:** Department of Anesthesia, UCSF Helen Diller Family Comprehensive Cancer Center, University of California at San Francisco, San Francisco, California 94110-1305

**Keywords:** Bispecific, Multi-specific, linker-enforced chain pairing, protease-aided linker removal, thrombin, LegoBody

## Abstract

Bispecific and multi-specific antibodies are capable of recognizing multiple ligands simultaneously or synergistically, creating complex biological interactions not achievable by monoclonal antibodies, thus expanding opportunities for novel therapy development. With the large number of monoclonal antibodies either approved or under clinical development, there are numerous opportunities to combine their specificities to further improve therapeutic potential. Although simple in concept, clinical development of bi- and multi-specific antibodies face several challenges, chief of which is how to efficiently and reliably produce bispecific and multi-specific antibodies with expected specificity and desired biophysical properties. In this study, we developed a modular approach that uses temporary linkers to enforce proper chain pairing and proteases such as thrombin to remove those linkers from the final product. Combined with the ‘knob-into-hole’ design, we can generate IgG-like, bi- or multi-specific antibodies from any pre-existing monoclonal antibodies. The approach is highly versatile and applicable to any monoclonal antibody pair or panel, expediting evaluation and therapeutic development of bi- and multi-specific antibodies.

## Introduction

Antibodies with bi- or multi-specificity hold great promise for the next generation of therapeutic drugs against a variety of diseases, including cancers, infections, and immunological disorders. Compared to traditional monoclonal antibodies that specifically recognize one ligand, bi- or multi-specific antibodies can recognize two or more ligands and thus may provide an advantage in co-engaging different cell types ^1-4^, creating synthetic specificity ^5^, altering internalization dynamics ^6, 7^, synergistically neutralizing virus and toxin ^8-10^, and simultaneously block multiple signaling pathways to maximize therapeutic benefits ^11-17^. In addition, bi- and multi-specific antibodies can increase sensitivity and breath for recognizing target cells, tissues, or pathogens ^8, 18-20^. Some bi-specific antibodies were also designed to possess desirable properties other than recognition, such as enhanced production, extended half-time, or increased tissue penetration ^21, 22^.

Production of bi- or multi-specific antibodies is more complicated than monoclonal antibodies^18, 23, 24^. Many different formats of bi-specific antibodies have been designed, including chemical conjugation of two disparate antibodies, tandem single-chain variable fragments (scFv) or Fab domains, and scFv or Fab fusion to immunoglobulins or other scaffold proteins ^18, 23, 25-34^. The design of multi-specific antibodies is often derived from these bispecific formats but with substantially higher complexity ^35-37^. Each of these designs, and their derivatives, features distinct topology and thus may have non-identical biological functions and different pharmacokinetics, which needs to be tested experimentally. Depending on the molecular form of the building block, some formats suffer from drawbacks, ranging from poor production, low *in vivo* stability, and immunogenicity ^23, 38^. Consequently, new methods and formats to produce bi- or multi-specific antibodies are important topics for exploration.

Among the various designs, the IgG-like format is our main area of interest due to its resemblance to natural antibody, which often have good yields, stability, and relatively low immunogenicity ^39, 40^. To produce these IgG-like molecules, two major problems have to be addressed: proper pairing of heavy chains from different antibodies, and correct pairing of the light chains to their corresponding heavy chains. The first issue is often addressed by the now classic knobs-into-holes (KIH) Fc mutants that enforce the formation of heavy chain heterodimers ^41, 42^. However, the desired heterodimer in such a design is not exclusive, with homodimers still present, likely due to insufficient thermal stability of the CH3 domain and the Fc interface of the KIH heterodimer ^43^. In fact, both knob and hole Fc domains can form homodimers, causing substantial contamination in products that were generated by the KIH approach alone ^44-48^. As the homodimer often shares similar biophysical features as the heterodimer, such as size and isoelectric point, optimization and purification of the KIH bispecific format can be challenging and time-consuming. Additional modifications of the KIH Fc have been explored to enhance the formation of heterodimer ^45-49^.

To avoid promiscuous association of light chains to heavy chains in bispecific antibody production, different antibodies (with different ligand binding specificity) using one common light chain have been selected from phage or yeast display libraries, providing a popular solution to the light chain pairing problem ^50-52^. However, this restriction on light chain reduces the sequence space that can be explored for binding affinity/specificity and downstream developability. The CrossMab format is an alternative approach for solving the light chain pairing problem, swapping only one pair of the variable or constant domains in one of the Fab fragments, and thus inhibiting chain mismatching between the switched Fab and the un-switched ones ^53^. Nonetheless, incorrect pairing of light chains are often observed, resulting in variable levels of product heterogeneity ^23^. The issue of immunogenicity of the hybrid variable-constant domain remains to be investigated.

The feasibility of single-chain IgG (scIgG) has been demonstrated by previous studies and can be used to resolve the light chain pairing problem ^54-56^. However, an extended linker between light and heavy chains, varying between 30-60 residues in length, was retained in the final product, limiting its utility. Here, we sought to develop an approach to produce single-chain IgG with cleavable linkers, allowing for removal of the undesired linkers *in vitro* through the use of proteases. We successfully introduced a thrombin cleavable linker between the light chain and the heavy chain, which can be removed by commercially available enzymes with high efficiency and accuracy. Antibodies produced using this approach show similar yield as the original antibodies, feature intact ligand-binding affinity, and contain only a few linker residues. Compared to an *in vivo* cleavage system designed in a previous study ^57^, the cleavage in our system happens post-translationally and post-purification, ensuring correct pairing of the chains during cleavage. Furthermore, the linkers enable us to engineer extra features in each of the two chains, such as different affinity tags, which can be used for easy purification of highly pure bispecific IgG molecules. Finally, unlike an approach that is designed to produce the entire bispecific IgG-like molecule as a single polypeptide before enzymatic processing ^58^, our approach is totally modular, demands far less on the protein synthesis machinery, and allows facile production of not only bispecific but also multi-specific antibodies, with IgG-like backbone and Fab-based binding modules.

## Results

### Monoclonal antibodies produced from thrombin-removable single-chain IgGs show similar biochemical features as native IgGs

Single-chain IgGs using peptide linkers to join the light chain and heavy chain have been investigated previously for production and ligand binding ^54-56^. The linker in scIgG remains a liability for therapeutic development. We therefore developed an efficient system to remove the peptide linker to obtain a true IgG-like molecule.

We tested several commercially available proteases and determined that thrombin is the most suitable enzyme because of its efficiency, accuracy, and compatibility with non-reducing environments. We chose the clinically established antibodies, Ipilimumab (Ipili) and Daratumumab (Dara) as the study antibodies and generated scIgGs of Ipili and Dara with a linker in-between the light chain and heavy chain. The linker, sc36TMB, is a flexible 36-residue peptide adapted from a previous study ^55^, with additional thrombin cleavage sites on both the N- and C-terminus (Fig.1A, Supplemental Fig. S1A).

**Fig. 1.**
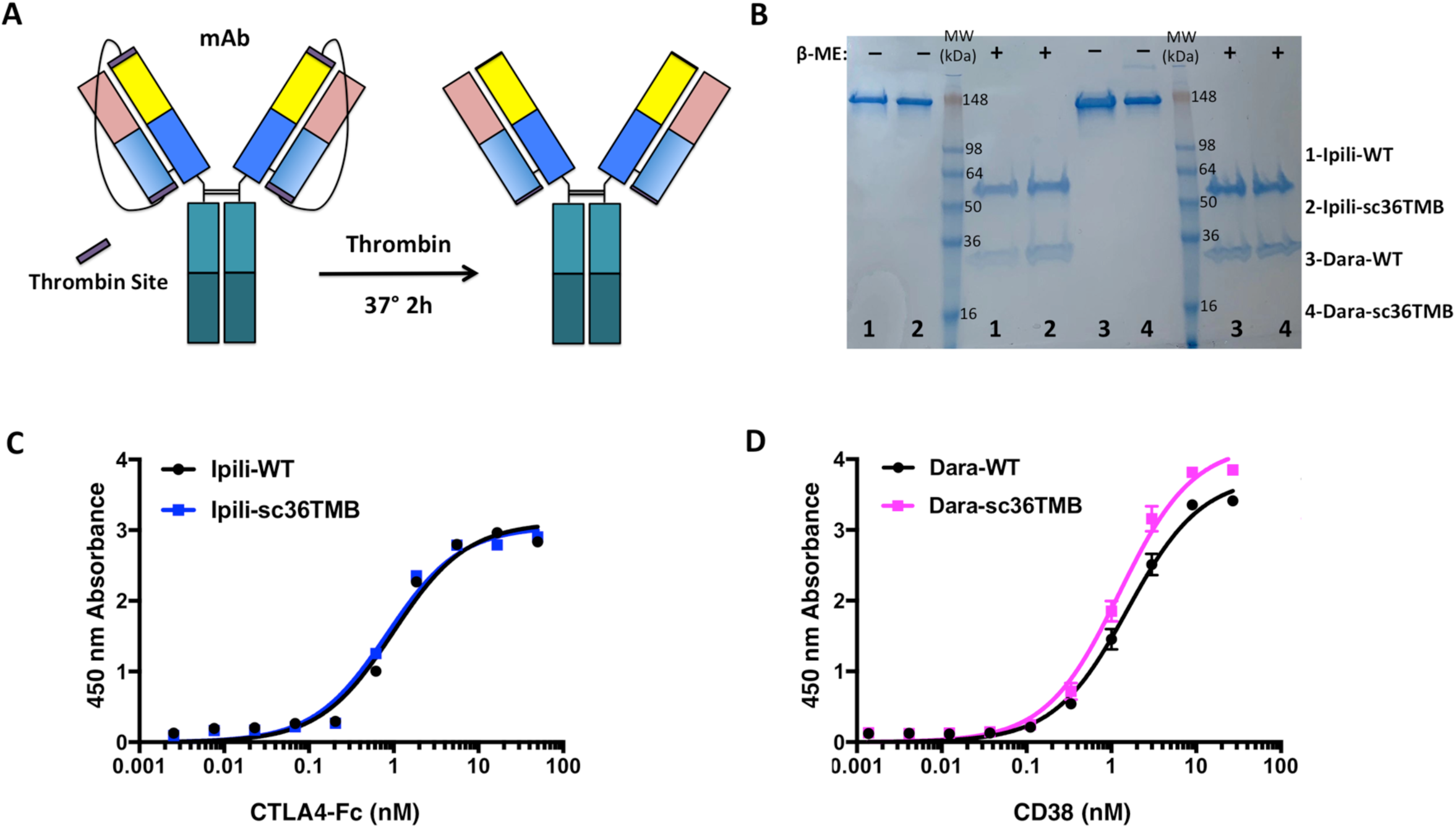
Monoclonal antibodies processed from single-chain IgGs show similar biophysical and binding properties as the original antibodies. (**A**) The scheme to produce monoclonal IgG from single-chain IgG with a thrombin cleavable linker. (**B**) After thrombin cleavage, the IgG molecules produced with the thrombin-cleavable linker (Ipili-sc36TMB and Dara-sc36TMB) show molecular weights of approximately 150 kDa and can be separated into heavy chains (∼50 kDa) and light chains (∼25 kDa) with a reducing agent, β-mercaptoethanol (β-ME), a virtually identical pattern compared with the original monoclonal antibody. MW: approximate molecular weight marker. (**C**) Ipili-sc36TMB and (**D**) Dara-sc36TMB bind to their respective ligands in ELISA assays, showing similar binding profiles compared with the original monoclonal antibodies.

The engineered scIgG displayed similar yield as the original antibodies (Supplemental Fig. S1B). Furthermore, the linker was successfully removed by thrombin cleavage, yielding a ∼150 kDa antibody product formed by disulfide-bonded light (∼25 kDa) and heavy chains (∼50 kDa), same as the natural antibody (Fig. 1B). Following thrombin cleavage, a 5-mer peptide (-GLVPR) remained at the C-terminus of the light chain, as well as two residues (-GS) at the N-terminus of the heavy chain (Supplemental Fig. S1A). To evaluate if the processed IgG molecule possesses an intact paratope, we tested ligand binding by ELISA (Fig. 1C, D). Ipilimumab binds to the CTLA4-Fc with an EC_50_ of approximately 0.99 ± 0.13 nM, while the EC_50_ of the cleaved Ipili-sc36TMB is estimated to be about 0.83 ± 0.11 nM (Fig. 1C). Likewise, Daratumumab binds its ligand, CD38, with EC_50_ of approximately 1.56 ± 0.11 nM, and the cleaved Dara-sc36TMB has an EC_50_ of 1.24 ± 0.09 nM (Fig. 1D). The similar EC_50_ values between the processed scIgGs and the original antibodies indicate that the engineered linker and the cleavage process do not interfere with the formation of the antibody or the integrity of its paratope.

### Bispecific antibodies produced from thrombin-cleavable single-chain IgGs with the ‘knobs-into-holes’ Fc heterodimer

We next expand the study from monoclonal to bispecific antibody generation. The KIH design features critically paired mutations in the ‘knob’ Fc (T366W) and ‘hole’ Fc (T366S/L368A/Y407V), which enforces pairing of the KIH Fc heterodimer ^41, 42^. However, this KIH design does not enforce proper light chain pairing.

The successful production and thrombin-cleavage of the scIgGs to IgG molecules provides a plausible solution for enforcing correct pairing between light and heavy chains. Thus, we sought to generate bispecific antibodies using the KIH Fc system in conjunction with a thrombin-removable linker. We chose two sets of antibodies, Ipilimumab pairing with Daratumumab, and Ipilimumab paring with Herceptin, as the test samples. Our scheme to produce bispecific antibodies is shown in Fig. 2A. In brief, the light chain of each antibody is fused with the corresponding heavy chain via the thrombin-removable linker. The Fc used for Ipilimumab is the ‘hole’ Fc, while the Fc for either Daratumumab or Herceptin is the ‘knob’ Fc. Bispecific antibodies were produced by co-transfecting HEK293A cells with the ‘knob’ and ‘hole’ scIgG constructs at a 1:1 ratio, and purified from culture supernatant by protein A agarose, and processed by thrombin to remove the linker.

**Fig. 2.**
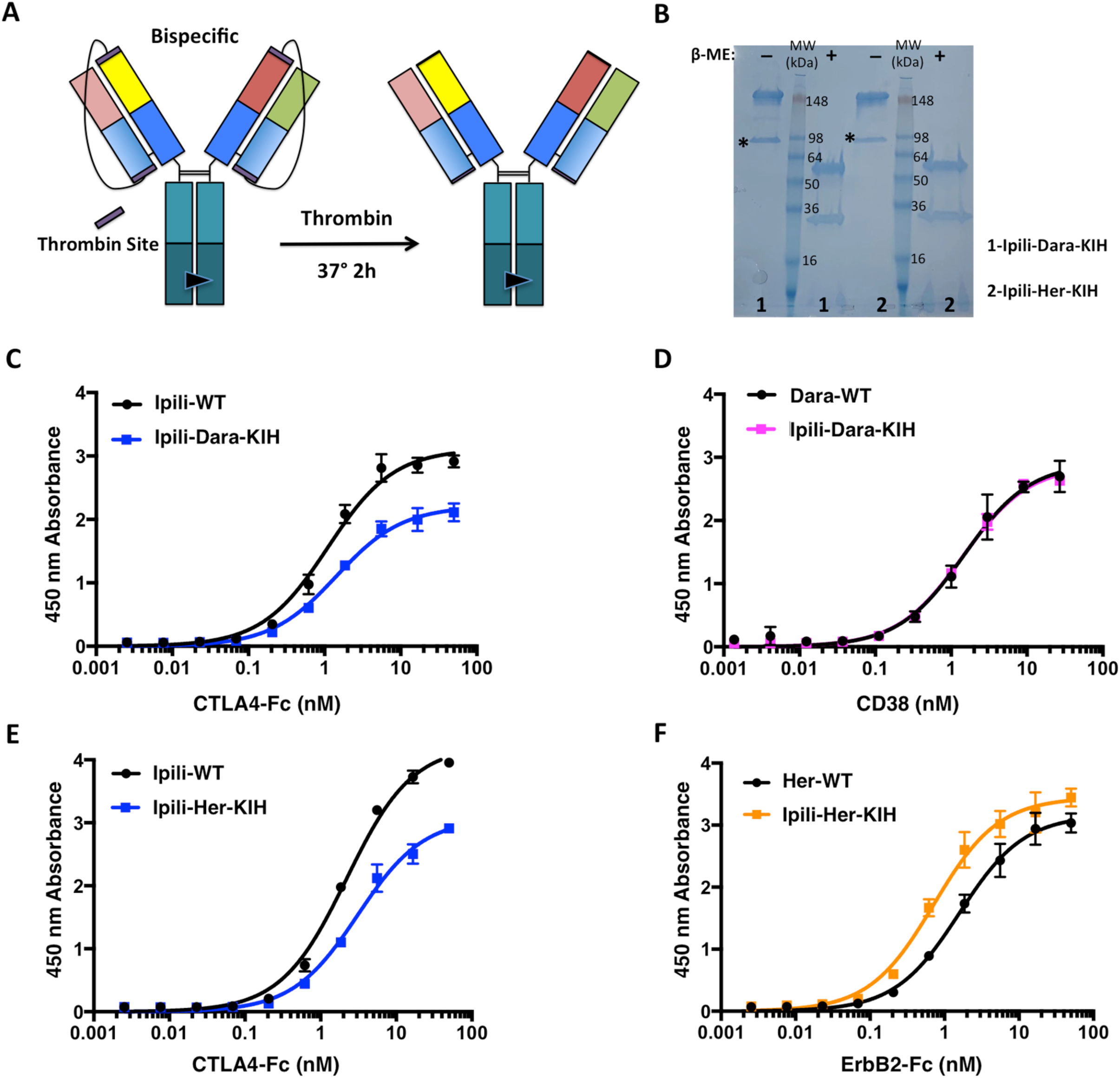
Bispecific antibodies produced with thrombin-cleavable linkers and KIH Fc domains. (**A**) The scheme to produce bispecific antibodies. (**B**) Two bispecific antibodies, Ipili-Dara-KIH and Ipili-Her-KIH, were generated, both of which show a predominant component with a molecular weight of 150 kDa. A small fraction of contamination, likely caused by incorrectly paired homodimers, can be observed at the half size of an antibody (labeled with ‘*’). Under reducing condition, separated heavy and light chains, ∼50 kDa and ∼25 kDa respectively, were observed. (**C, D**) Bispecific Ipili-Dara-KIH binding to CTLA4-Fc (**C**) and CD38 (**D**), in comparison to the original monoclonal antibodies. (**E, F**) Bispecific Ipili-Her-KIH binding to CTLA4-Fc (**E**) and ErbB2-Fc (**F**), in comparison to the original monoclonal antibodies.

Using this system, we were able to obtain relatively pure IgG-like molecules for both bispecific antibodies with estimated molecular weights of ∼150 kDa under non-reducing condition (Fig. 2B). The KIH product can be separated into light chain (∼25 kDa) and heavy chain (∼50 kDa) by reduction with β-mercaptoethanol (β-ME) (Fig. 2B). We assessed the purity of the KIH product using analytical hydrophobic interaction chromatography (HIC). Both Ipili-Dara-KIH and Ipili-Her-KIH display a dominant bispecific component, comprising 71% and 73% of the total product, respectively (Fig. S2, S3).

We also investigated the integrity of the paratopes of each Fab in the bispecific by ELISA. Unlike an IgG molecule that binds bivalently to one ligand, the binding of the IgG-like bispecific to each of its two ligands is monovalent. To minimize the influence of valency in the ELISA assay, we mobilized antibodies on plates and assessed binding to ligands in solution. As shown in Fig. 2C & D, and Supplemental Fig. S4A, the Ipili-Dara-KIH is capable of binding to both CTLA4 and CD38, in manners that are similar to the parental monoclonal antibodies (Ipilimumab and Daratumumab), indicating that the bispecific antibodies possess intact paratopes. Similar results were observed for the bispecific Ipili-Her-KIH (Fig. 2E & F, and Supplemental Fig. S4B).

### Use of linker-embedded affinity tags for further purification of bispecific antibodies

Contamination in bispecific antibodies produced by the KIH approach has been observed previously ^45-47, 49^, which primarily arises from ‘hole-hole’ or ‘knob-knob’ homodimers. Common procedures for antibody purification, such as protein A or G affinity chromatography, reply on recognition of the Fc domain and thus cannot effectively distinguish the bispecific KIH heterodimer from contaminating homodimers. Prior approaches to enhance the efficiency of correct pairing of the KIH heterodimer or eliminate the non-bispecific contamination often involve introducing additional mutations and additional steps in purification.

Here, we took advantage of the removable nature of the linker and embedded a pair of affinity tags, the 10-His tag and Twin-Strep-Tag^®^, into the linkers to purify the bispecific product (Fig. 3A). The KIH heterodimer would possess both affinity tags and thus be readily distinguished from homodimers. We constructed plasmids expressing Ipilimumab with a Twin-Strep-Tag^®^ linker and the ‘hole’ Fc mutant, and Daratumumab or Herceptin with a 10-His tag linker and the ‘knob’ Fc mutant, co-transfected HEK293A cells to generate bispecific antibodies, i.e., Ipili-Dara-KIH and Ipili-Her-KIH. The supernatant was sequentially purified using Ni-NTA and Strep-Tactin^®^XT resin to obtain antibody products with both tags and thus both specificities. The linkers were removed by thrombin cleavage to generate the final product. SDS-PAGE shows that bispecific antibodies produced in this manner contain substantially lower amounts of containments (labeled with ‘*’) (Fig. 3B). HIC-HPLC analysis showed that the purities of the Ipili-Dara-KIH and Ipili-Her-KIH are nearly 100% and about 92%, respectively (Fig. 3C, D).

**Fig. 3.**
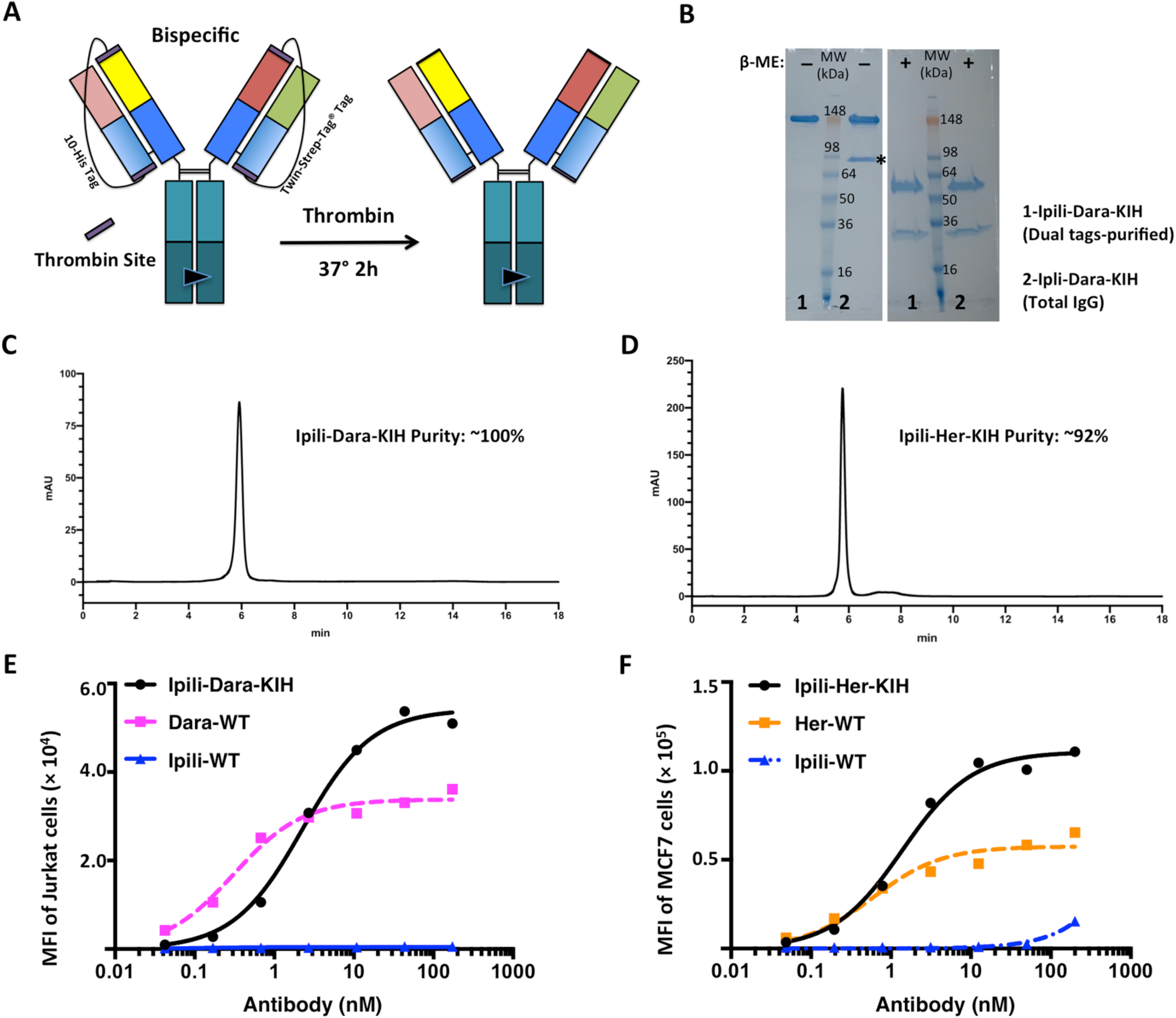
Bispecific antibody with high purity produced with a pair of thrombin-cleavable dual-tagged affinity linkers. (**A**) The scheme to produce bispecific antibodies with built-in affinity tags, e.g., 10-His tag and Twin-Strep-Tag^®^ that enables sequential purification of the desired heterodimer by Ni-NTA resin and Strep-Tactin^®^XT resin. (**B**) The bispecific product of Ipili-Dara-KIH purified via the dual tags (Ni-NTA and Strep-Tactin^®^XT) is compared with the same product purified by protein A agarose. The contamination in the bispecific products that can be observed at the half-antibody size (the ‘*’ band) is eliminated when purified with dual tags. (**C, D**) The purity of bispecific products is assessed by analytical HIC. (**E**) Flow cytometry analysis of Jurkat cell binding by Ipili-Dara-KIH, Daratumumab, and Ipilimumab at various concentrations (0 - 200 nM). (**F**) Flow cytometry analysis of MCF7 cell binding by Ipili-Her-KIH, Herceptin, and Ipilimumab at various concentrations (0 - 200 nM).

We next studied how bispecific antibodies recognized cells expressing target antigens. The Jurkat cell line has been used to study CD38 binding ^59, 60^. We found that indeed Jurkat cells express CD38 and are bound by Daratumumab (Fig. 3E). However, little to no expression of CTLA4 was detected by Ipilimumab (Fig. 3E). Thus Jurkat cells are useful mainly for assessing the CD38 binding arm. The bispecific antibody, Ipili-Dara-KIH, which is monovalent in each specificity, binds to Jurkat cells with EC_50_ of 2.26 ± 0.26 nM (Supplemental Fig. S5A). Daratumumab, a bivalent IgG, binds to Jurkat cells with EC_50_ of 0.31 ± 0.05 nM. The bispecific Ipili-Dara-KIH binds to Jurkat cells with a higher median fluorescence intensity (MFI) value compared with Daratumumab, consistent with its monovalent binding mode.

Similarly, we evaluated how the bispecific Ipili-Her-KIH antibody binds to the breast cancer cell line MCF7 that has been reported to express both ErbB2 (Her2) and CTLA4 ^61^. We observed a high staining signal by flow cytometry on MCF7 by Herceptin, but a rather low signal by Ipilimumab. Thus we used MCF7 cells to evaluate the Her2-binding arm only. The bispecific Ipili-Her-KIH binds to MCF7 (Fig. 3F) with EC_50_ of 1.39 ± 0.21 nM (Supplemental Fig. S5A). Herceptin, a bivalent IgG, binds to MCF7 (Fig. 3F) with EC_50_ of 0.60 ± 0.18 nM (Supplemental Fig. S5A). The total MFI for the bispecific Ipili-Her-KIH is higher than Herceptin, consistent with its monovalent binding mode.

To evaluate binding of bispecific antibodies to CTLA4 on cell surface, HEK293A cells were transiently transfected with a construct expressing the human *CTLA4* gene. After culturing for 18 hours, the cells were harvest and analyzed by flow cytometry for binding by Ipilimumab IgG, Ipilimumab Fab, Ipili-Dara-KIH, and Ipili-Her-KIH. Like Ipilimumab, both bispecific antibodies were able to bind to the CTLA4-transfected cells (Supplemental Fig. S5B), confirming that the anti-CTLA-4 arm is functional. The apparent binding affinities of the bispecific antibodies are lower than that of the bivalent Ipilimumab IgG, an expected result given the valency difference, but higher than the Ipilimumab Fab that is also monovalent (Supplemental Fig. S5C). MFI values at saturation binding are higher for the bispecific (and the Ipilimumab Fab) than the Ipilimumab IgG, again consistent with the monovalent vs. bivalent binding mode.

### Facile generation of multi-specific antibodies using thrombin-removable linkers

The generation and application of antibodies with even higher order of specificity and complexity are challenging tasks that have only been attempted a limited number of times ^35-37^. In general, as the complexity of the molecule grows, the difficulty grows disproportionally or exponentially in proper assembly of six, eight, or more chains into one antibody. Here, we take advantage of the modular nature of the linker-enforced chain assembly and generate multi-specific antibodies using the enzymatically cleavable linker. In simplified terms, these molecules were designed based on the IgG-like bispecific with additional specificities introduced by appending Fab domains to either the N-terminus (Fig. 4 & 5, tandem Fabs) or C-terminus (Fig. 6) of the bispecific molecule.

**Fig. 4.**
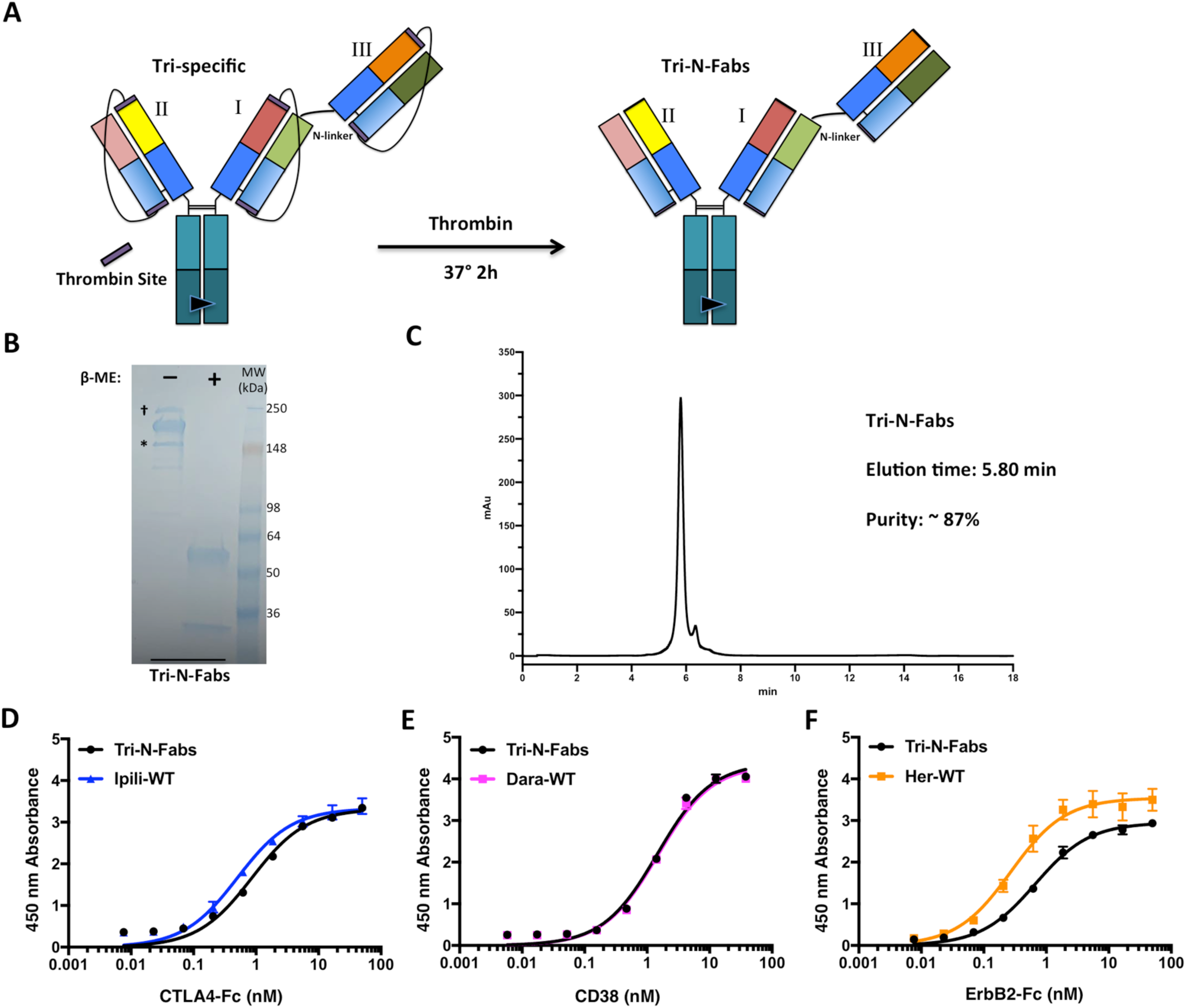
Design of Tri-N-Fabs as a new format for tri-specific antibodies produced with thrombin-removable linkers and KIH Fc heterodimers. The production scheme and final structure of the Tri-N-Fabs is presented in (**A**). The extra Fab domain (III) is appended to the N-terminus of the bispecific antibody. For the Tri-N-Fabs produced, we placed Fabs of Ipilimumab at position I, Daratumumab at position II, and Herceptin at position III. (**B**) On SDS-PAGE, the Tri-N-Fabs shows an approximate molecular weight of 200 kDa and can be separated into polypeptide chains of ∼50 kDa and ∼25 kDa under reducing conditions. Small fractions of contamination were observed, marked as (*) and (†), likely caused by homodimers. (**C**) The purity of the Tri-N-Fabs is assessed by analytical hydrophobic interaction chromatography. (**D, E, F**) Tri-N-Fabs interacts with the three intended ligands in an ELISA assay. Parental antibodies were included as references.

**Fig. 5.**
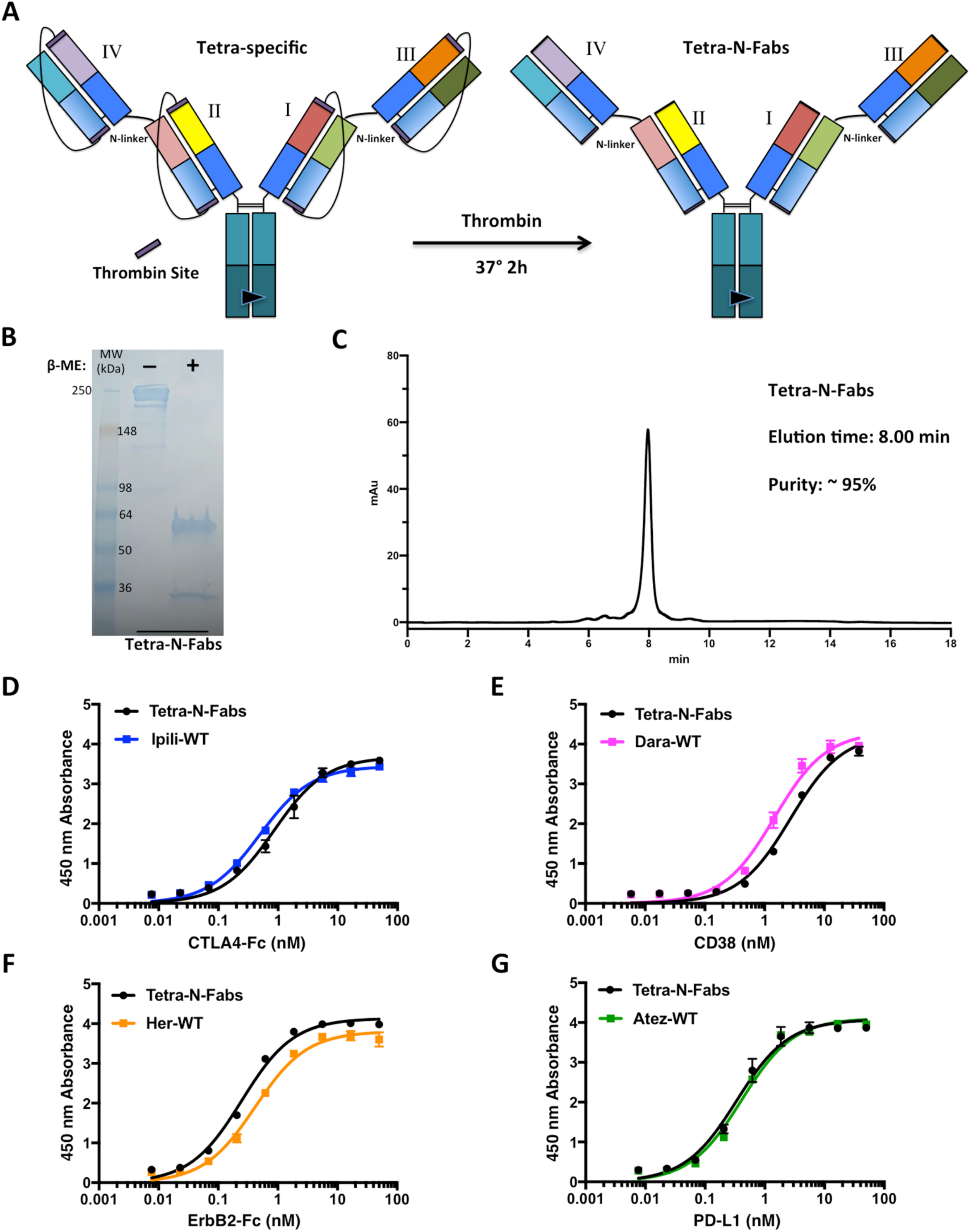
Tetra-N-Fabs, a new format for tetra-specific antibodies, is produced with thrombin-removable linkers and KIH Fc heterodimers. (**A**) The design of Tetra-N-Fabs. Extra Fab domains (III and IV) are appended to the N-terminus of the bispecific antibody. For the Tetra-N-Fabs produced, we placed Fabs of Ipilimumab at position I, Daratumumab at position II, Herceptin at position III, and Atezolizumab at position IV. (**B**) The Tetra-N-Fabs displays a molecular weight of ∼250 kDa by non-reducing SDS-PAGE analysis and can be separated into two polypeptide chains of ∼50 kDa and ∼25 kDa under reducing conditions. (**C**) The purity of the Tetra-N-Fabs is estimated to be 95% by analytical HIC. (**D, E, F, G**) Binding of the Tetra-N-Fabs to all four ligands with parental antibodies included as references.

**Fig. 6.**
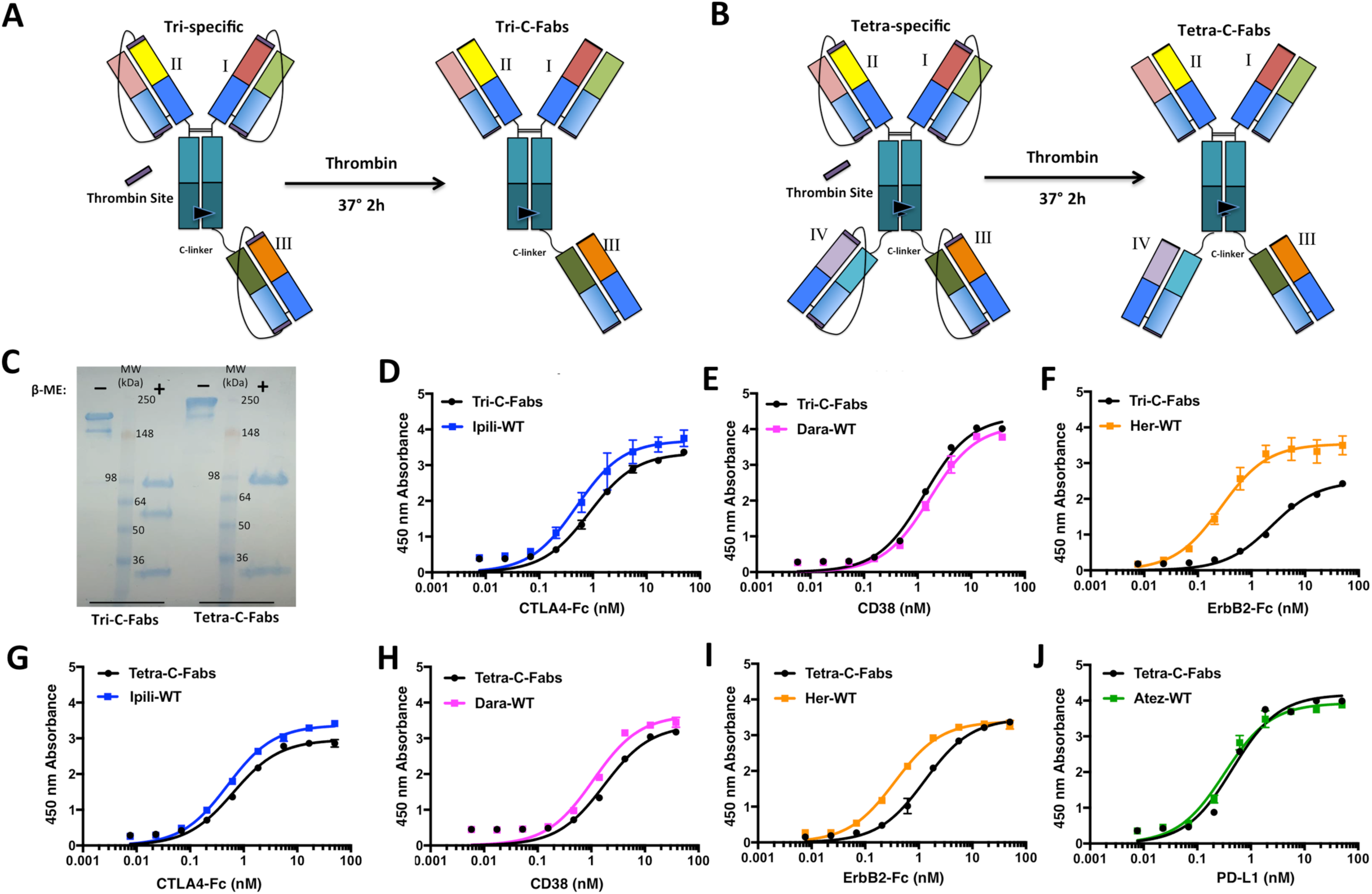
Tri-C-Fabs and Tetra-C-Fabs are alternative formats for tri- and tetra-specific antibody generation using the thrombin-removable linker and KIH Fc heterodimer. Schematic presentations of Tri-C-Fabs (**A**) and Tetra-C-Fabs (**B**). Fab domains representing the 3^rd^ and 4^th^ specificity were appended to the C-terminus of the bispecific antibody. We placed Fabs of Ipilimumab at position I, Daratumumab at position II, Herceptin at position III, and Atezolizumab at position IV (in the case of Tetra-C-Fabs). (**C**) SDS-PAGE analysis of Tri-C-Fabs and Tetra-C-Fabs. The Tri-C-Fabs displays an approximate molecular weight of 200 kDa and can be separated into polypeptide chains of ∼75 kDa, ∼50 kDa, and ∼25 kDa under reducing conditions. The Tetra-C-Fabs shows an approximate molecular weight of 250 kDa and can be separated into polypeptide chains of ∼75 kDa and ∼25 kDa under reducing conditions. (**D, E, F**) The Tri-C-Fabs binds to all three intended ligands by ELISA, with the parental antibodies included as references. (**G, H, I, J**) Binding of the Tetra-C-Fabs to all four intended ligands by ELISA, with parental antibodies included as references.

In the case of tandem Fab constructs shown in Fig. 4 (tri-specific) and 5 (tetra-specific), the four different chains (two heavy and two light chains) of two antibodies (I, III) are genetically fused into one polypeptide in the following configuration: light chain−sc36TMB linker−heavy chain−N-linker−light chain (I)−sc36TMB linker−heavy chain (I) and ‘hole’ Fc mutant (shown in Fig. 4A). This tandem Fab construct is paired with one antibody construct (position II) with the ‘knob’ Fc mutant to produce the tri-specific antibodies (Fig. 4A). To produce the tetra-specific molecule, another two antibodies (II, IV) were constructed in a second tandem Fab construct with the ‘knob’ mutant in a configuration similar to that of the ‘hole’ Fc mutant (Fig. 5A).

Co-expression of corresponding constructs produces the precursors for either tri-specific (Fig. 4A) or tetra-specific (Fig. 5A) molecules, which are processed by thrombin cleavage to remove the linker scaffold within each Fab unit. In this way, we were able to generate the asymmetric tri-specific antibody (Tri-N-Fabs) and the symmetric tetra-specific antibody (Tetra-N-Fabs), with Ipilimumab in position I, Daratumumab in position II, Herceptin in position III, and Atezolizumab in position IV (in the case of Tetra-N-Fabs).

Purified Tri-N-Fabs and Tetra-N-Fabs displayed correct molecular weights by SDS-PAGE analysis, estimated to be 200 kDa and 250 kDa respectively, under non-reducing conditions and were separated into polypeptides with molecular weights of approximately 50 kDa and 25 kDa under reducing conditions (Fig. 4B & 5B). Purity of the final products was assessed by analytical HIC-HPLC and found to be approximately 87% for Tri-N-Fabs (Fig. 4C) and 95% for Tetra-N-Fabs (Fig. 5C). We evaluated antigen binding by ELISA, where the antibody was immobilized on plate, and tested for binding to biotinylated ligands in solution (CTLA4-Fc, CD38, ErbB2-Fc, and PD-L1). Both the Tri-N-Fabs and the Tetra-N-Fabs showed binding to all corresponding ligands with apparent affinity similar to that of the parental antibodies (Fig. 4D-F and Supplemental Fig. S7A for Tri-N-Fabs; Fig. & 5D-G and Supplemental Fig. S7B for Tetra-N-Fabs). These results indicate that the Tri-N-Fabs and the Tetra-N-Fabs possess the designed paratopes that are functional.

An alternative way to arrange the different Fab modules in a tri- or tetra-specific antibody is to append the extra Fabs to the C-terminus of the Fc domain (Fig. 6A for the tri-specific, Tri-C-Fabs; and Fig. 6B for the tetra-specific, Tetra-C-Fabs). For the asymmetric Tri-C-Fabs, the two chains are in the following configuration: light chain (I)−sc36TMB linker−Fab heavy chain (I)−‘hole’ Fc mutant−C-linker−light chain (III)−sc36TMB linker−Fab heavy chain (III); and light chain (II)−sc36TMB linker−Fab heavy chain (II)−‘knob’ Fc mutant (Fig 6A). The C-linker was designed as a 13-residue long Gly-Ser peptide. For the symmetric Tetra-C-Fabs, light chain (I/II)−sc36TMB linker−Fab heavy chain (I/II)−‘hole’/‘knob’ Fc mutant−C-linker−light chain (III/IV)−sc36TMB linker−Fab heavy chain (III/IV) (Fig. 6B). These multi-specific antibodies were produced by co-transfecting plasmids expressing the corresponding chains followed by purification on protein A agarose. The linkers in-between light and heavy chains were removed by thrombin cleavage prior to analysis.

We successfully produced a tri-specific antibody (Tri-C-Fabs) with Ipilimumab Fab at position I, Daratumumab Fab at II, and Herceptin Fab at III; and a tetra-specific antibody (Tetra-C-Fabs) with the addition of Atezolizumab Fab at IV. The main products for both Tri-C-Fabs and Tetra-C-Fabs displayed the correct molecular weights by SDS-PAGE analysis (Fig. 6C). Analytical HIC-HPLC indicated that the overall purities of the Tri-C-Fabs and Tetra-C-Fabs were approximately 93% and 79%, respectively (Supplemental Fig. S6A, B). Both Tri-C-Fabs and Tetra-C-Fabs were able to bind their intended ligands by ELISA, with EC_50_ similar to that of the parental antibodies, i.e., Ipilimumab, Daratumumab, and Atezolizumab (Fig. 6D-J, Supplemental Fig. S7C, D). We observed a reduction in target binding by the Herceptin Fab in these multi-specific products. With regard to ErbB2 binding, the Tri-C-Fabs and Tetra-C-Fabs display approximately 9- and 3-fold higher EC_50_ values compared to the parental antibody Herceptin. However, the effect seems to be unique to the Herceptin Fab as the Atezolizumab Fab in a similar position in Tetra-C-Fabs was not affected (Fig. 6J).

We also evaluated the binding kinetics of the bispecific or multi-specific antibodies to corresponding ligands by biolayer-interferometry (BLI). We immobilized the biotinylated ligands on the streptavidin-coated sensor and measured antibody binding. We found that the bispecific and multi-specific antibodies bind to their ligands in manners expected from their monovalent binding mode (Fig. S8). The parent monoclonal IgGs showed lower K_d_ values due to avidity (bivalent vs. monovalent), but the parent monovalent Fab (Ipilimumab Fab) showed similar K_d_ as the bipsecific and multi-specific antibodies.

## Discussion

In this study, we have developed a facile and modular platform to generate bispecific and multi-specific antibodies from any pre-existing monoclonal antibody. The use of thrombin-removable linkers enforces correct pairing of light and heavy chains and enables efficient *in vitro* removal of these linkers following purification. Comparing other bispecific formats, such as the CrossMAb or common light chain IgG molecules, our format requires minimal efforts in design and optimization, imposes no restrictions on the underlying monoclonal antibody (the building block), and can be used as tools to readily generate and evaluate bispecific antibodies from any two monoclonal antibodies of interest. The linker we introduced to enforce proper chain paring is very malleable and can be used to introduce versatile features to aid purification. As a proof of concept, we have designed different affinity tags into the linker, and demonstrated that sequential purification by affinity chromatography can effectively eliminate homodimer contaminations. Other features such as asymmetric length or charge (isoelectric point) can be incorporated into the linker to customize the purification scheme.

We further expanded the approach, which is modular in nature and permits Lego-like assembly, to multi-specific antibody generation. We generated two classes of tri-specific and tetra-specific antibodies by appending additional Fab domains to either the N-terminus of the light chain or the C-terminus of the heavy chain in the starting bispecific molecule. The added Fab domains also use thrombin-removable linkers to enforce correct heavy and light chain pairing. We have successfully generated several tri- and tetra-specific molecules and verified their binding to corresponding ligands. Given the modular nature of the process, our approach should be applicable to the generation of even higher order of specificities. This marks a major distinction from an approach where the IgG-like bispecific is produced as a single giant polypeptide chain before enzymatic processing ^58^, which has production issues due to the sheer size of the molecule and is essentially limited to bispecific only. Given the explosion of information on the molecular mechanism of human diseases and the rapid expansion of the list of potential targets, multi-specific antibodies are likely to become an expanding class of novel therapeutics due to their unique ability to generate synergistic or synthetic interactions among targets, thus uncovering new biology and targeting opportunities. Our approach offers a means of facile generation of novel molecular tools to fully explore those opportunities.

## Materials and Methods

### Monoclonal antibody expression and purification

Genes encoding antibody variable domains were synthesized by gBlock^®^ (Integrated DNA Technologies). Plasmids for the heavy and light chains for each antibody were separately cloned in the Abvec vector as previously described ^5, 62^. Antibodies were produced by co-transfecting plasmids expressing the heavy and light chains at a 1:1 ratio in HEK293A cells for 6-8 days, followed by purification of culture supernatant on protein A agarose (Pierce/Thermo Scientific).

### Monoclonal single-chain IgG expression and purification

The sc36TMB linker (*GLVPRGSGSGGGSGGGSEGGGSEGGGSEGGGSEGGGSGGGSGGLVPRGS*), and the heavy chain variable domain were fused by PCR reaction. The resultant DNA fragment was subcloned into Ig-γ Abvec vector and fused with the heavy chain constant domain and Fc fragment to generate the complete single-chain IgG. The plasmid was transfected into HEK293A cells and scIgG molecules were purified from culture supernatant by protein A agarose (Pierce/Thermo Scientific).

### Thrombin cleavage and removal of the enzyme

To remove the sc36TMB linker, scIgG was mixed with thrombin at the ratio of 50-100 µg antibody per unit of thrombin (Millipore, 605160), and the mixture was incubated at 37°C for 2 hours. The processed IgG molecule was re-purified by protein A agarose to remove the enzyme. The conversion of the scIgG to IgG was verified by SDS-PAGE with reducing agent added to the sample.

### Bispecific antibody expression and purification

The ‘knob’ Fc (T366W) or ‘hole’ Fc (T366S/L368A/Y407V) fragment ^41, 42^ were introduced to the Ig-γ chain by site-directed mutagenesis. The fused gene encoding the complete light chain, sc36TMB linker, and heavy chain variable domain was subcloned into either the ‘knob’ or ‘hole’ vector as indicated in Results. To produce the scIgG bispecific antibody, a pair of the ‘knob’ and ‘hole’ vectors, each encoding one scIgG antibody, were co-transfected into HEK293A cells at a 1:1 ratio. After culturing for 6-8 days, the bispecific scIgG was purified from culture supernatant by protein A agarose. The sc36TMB linkers in the bispecific scIgG were removed by thrombin cleavage as described above, yielding the bispecific IgG.

### Tri-specific and Tetra-specific antibody expression and purification

The *AgeI* site between the signal peptide and N-terminus of the antibody gene in the Abvec vector was retrained in the construct for scIgG with either the ‘knob’ or ‘hole’ Fc. This *AgeI* site was used to introduce an additional Fab gene (with a thrombin removable linker in-between the light and heavy chain) and the N-linker (-ASTKGPSGSG-) by ligase independent cloning ^63^. To produce the tri- and tetra-specific antibodies, we co-transfected the pair of ‘knob’ and ‘hole’ vectorinto HEK293A cells, collected supernatant after 6-8 days which was purified on protein A agarose, and removed the sc36TMB linkers by thrombin. For producing the tri-specific antibody, the tandem Fab (Ipilimumab and Herceptin) with the ‘hole’ Fc vector was paired the single-chain Daratumumab vector with ‘knob’ Fc. For the tetra-specific, the tandem Fab (Ipilimumab and Herceptin) with the ‘hole’ Fc vector was paired the tandem Fab (Daratumumab and Atezolizumab) with ‘knob’ Fc vector. Both Tri-N-Fabs and Tetra-N-Fabs molecules were submitted for ion-exchange chromatography for additional purification.

Alternatively, a *HindIII* site at the C-terminus of the antibody gene was used to introduce the C-linker (-GGGSGGGSGGGSG-) and an extra Fab domain to the ‘knob’ or ‘hole’ vector. To produce the Tri-C-Fabs molecule, the ‘hole’ vector with two Fab modules (Ipilimumab at N-terminus and Herceptin at C-terminus) was co-expressed with the ‘knob’ Fc vector for the single-chain Daratumumab Fab in HEK293A cells. To produce the Tetra-C-Fabs molecule, the above ‘hole’ vector was co-expressed with the ‘knob’ Fc vector with two Fab modules (Daratumumab at N-terminus and Atezolizumab at C-terminus) in HEK293A cells. Following purification by protein A agarose, the unnecessary linkers were removed by thrombin. Ion-exchange chromatography was applied to improve the purity.

### Purification by ion-exchange chromatography

The tri-specific and tetra-specific molecules were further purified by ion-exchange chromatography after the removal of intra-Fab linkers. In brief, the molecules were applied to a Mono S™ 5/50 GL column (GE Healthcare) on ÄKTA (GE Healthcare). The following buffers were used: mobile phase buffer A: 20 mM MES (2-morpholin-4-ylethanesulfonic acid), pH 6.0; and mobile phase buffer B: 20 mM MES, 1 M NaCl, pH 6.0. After loading and washing the samples with buffer A, gradient elution with 10% to 50% B were used for fractionation. Fractions were collected and analyzed by SDS-PAGE.

### Ligand preparation and biotinylation

Extracellular domain Fc fusions for human CTLA4 (CTLA4-Fc) and ErbB2 (ErbB2-Fc) were purchased from Abcam (ab180054 and ab168896). These ligands were biotinylated by the EZ-link^®^ Sulfo-NHS-LC-Biotin according to the manufacturer’s protocol (Thermo Scientific). Excessive biotin was removed by buffer-exchange into PBS, and biotinylated ligands were concentrated and stored at −80°C. The recombinant extracellular domain of CD38 and PD-L1 was produced as a 6-His and AviTag™ fusion in HEK293 cells and purified by Ni-NTA followed by *in vitro* biotinylation by BirA biotin ligase (Avidity).

### ELISA and EC_50_ estimation

To accommodate the differences in valency between monoclonal and bi-, tri-, or tetra-specific antibodies, we performed the ELISA assay by immobilizing the antibody on a microtiter plate and testing binding to ligands in solution. The antibodies were diluted to 1 µg/ml in PBS and 100 µl of this solution per well were applied to the Nunc MaxiSorp ELISA plate (Thermo Fisher) for coating overnight. The plate was washed three times with PBS, blocked with 4% BSA at RT for 1 hour, and incubated with corresponding biotinylated ligands at various concentrations at RT for 1 hour, each condition in triplicates. The plate was then washed five times with the washing buffer (0.05% Tween-20 in PBS), incubated with 0.1 µg/ml HRP-conjugated streptavidin (Pierce/Thermo Fisher) at RT for 30 min, washed 3 times with the washing buffer, and incubated with the peroxidase substrate solution (SureBlue^®^, Seracare) at RT for 3-5 min before the reaction was terminated by adding equal volume of 1 M HCl. The absorbance at 450 nm of each well was detected by a microtiter plate reader (Synergy BioTek^®^ instrument). The absorbance value as a function of the ligand concentrations was analyzed to obtain the EC_50_ value by curve fit (Prism, GraphPad).

### Purity assessment by analytical hydrophobic interaction chromatography

Purified antibodies were analyzed by HIC-HPLC with the infinity 1220 LC System (Agilent). Mobile phase A consisted of 25 mM phosphate and 1.5 M ammonium sulfate, pH 7.0; and mobile phase B 25% (v/v) isopropanol in 25 mM phosphate, pH 7.0. Antibody samples were loaded onto a TSKgel HIC column (Tosoh Bioscience) and operated at 0.5 mL/min with a gradient from 10% to 100% B. Absorbance was detected at 280 nm. Purity was estimated by area integration using OpenLab CDS software (Agilent).

### Cell binding analysis by flow cytometry

Jurkat and MCF7 cell lines were obtained from American Type Culture Collection (ATCC) and cultured in Dulbecco’s Modified Eagle Medium (DMEM) with 10% fetal calf serum and 100 µg/ml penicillin-streptomycin in humidified atmosphere of 95% air and 5% CO_2_ at 37 ^°^C. For flow cytometry analysis, approximately 50,000 cells were incubated with different monoclonal, bispecific or multi-specific antibodies (the highest concentration of 200 nM with serial 4-fold dilutions) at RT for 1 hour, washed 3 times with PBS and further incubated with the secondary antibody solution (Alexa Fluor 647^®^-conjugated goat anti-human IgG, final concentration of 1 µg/ml) at RT for 1 hour, washed 3 times with PBS and analyzed by flow cytometry (Accuri™C6, BD Biosciences). The median fluorescence intensity (MFI) values were analyzed by Prism (GraphPad) to obtain EC_50_ values by curve fitting.

### K_d_ determination by bio-layer interferometry

A Gator instrument (Probe Life) was used to determine K_d_ of antibody-ligand interactions by bio-layer interferometry. Biotinylated ligands were diluted to 5-10 µg/ml in the provided kinetics buffer (Probe Life) and immobilized onto streptavidin-coated biosensors. The sensors were sequentially incubated with antibodies (200 nM) and kinetics buffers for 180 seconds to assess rates of association and disassociation. K_d_ values were analyzed by the Gator software (Probe Life).

## Author contributions

S.L. and B.L designed the project and wrote the manuscript. S.L. performed overall experiments. S.Y. performed analytical HPLC and provided anti-Erbb2 and CD38 antibodies. S.B. helped with FACS experiments and provided knob and hole vectors. N.L. provided PD-L1 ligands. BL conceived the overall project idea. All authors edited the manuscript.

## Acknowledgement

We thank Weihua Wen for help with protein purification. This work is supported in part by grants from the National Institutes of Health (R01 CA118919, R01 CA129491, R01 CA171315 and R01 CA223767).

## Declare of Interest

The authors are inventors of a patent related to this work.

