## Supplemental materials for "LegoBody: facile generation of bispecific and multi-specific antibodies"

**A** Single-chain IgG sequence:

(sc36TMB linker)

(Light chain)-*GLVPRGSGSGGGSGGGSEGGGSEGGGSEGGGSEGGGSGGGSGGLVPRGS*-(heavy chain)

Thrombin → 37° 2h

(Light chain)-GLVPR + GS-(heavy chain)

**B**

| General Production | HEK293A/100mm-dish/4-Day expression |
|---|---|
| Ipilimumab | 26μg |
| Ipli-sc36TMB | 28 μg |
| Daratumumab | 32μg |
| Dara-sc36TMB | 36 μg |

**Fig. S1 Design and production of single-chain IgG with thrombin cleavable linker.** (**A**) Sequence of the sc36TMB linker with the thrombin cleavage site marked with arrows. (**B**) The estimated yield of scIgG is comparable to that of the original antibody. Antibodies were purified by protein A affinity capture from culture supernatant collected from cell culture dish (100 mm in diameter) 4 days following transient transfection of HEK293A cells.

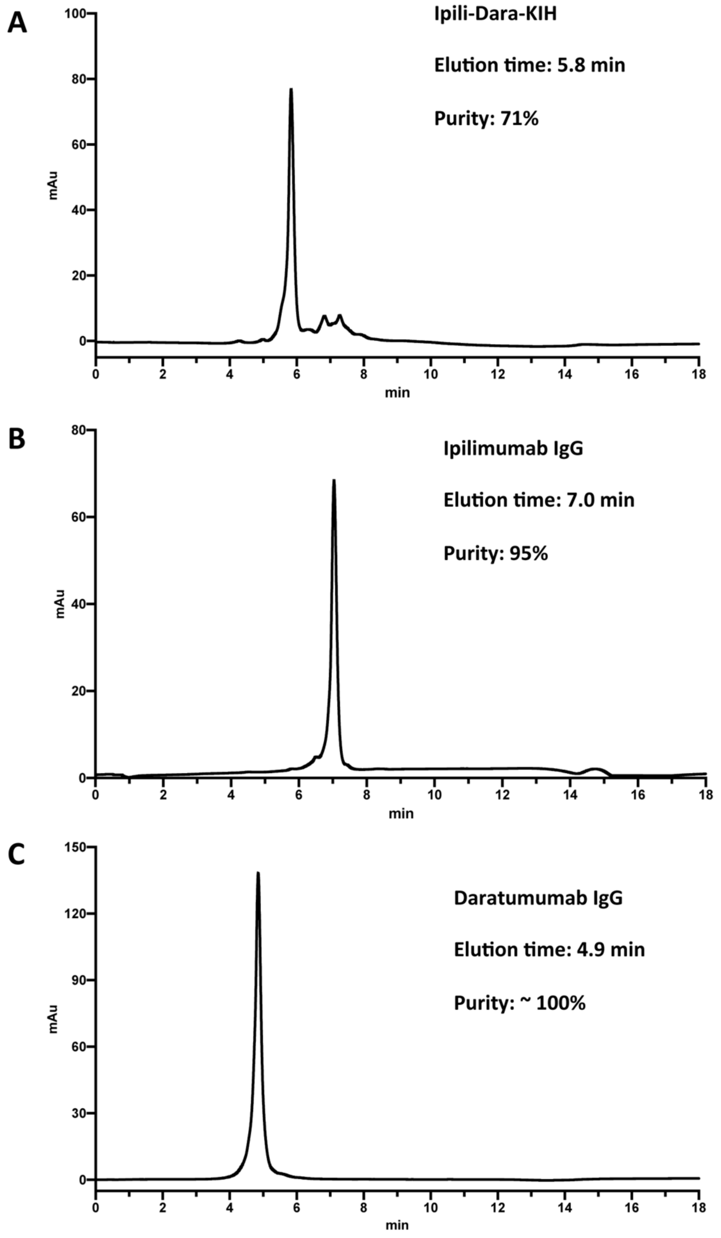


**Fig. S2 Purity assessment of the bi-specific antibody Ipili-Dara-KIH by analytical hydrophobic interaction chromatography.** The main peak of Ipili_Dara_KIH (**A**) shows an elution time in-between of Ipilimumab (**B**) and Daratumumab (**C**), representing the desired bispecific product that is estimated by area integration (OpenLab CDS, Agilent) to be 71% of total protein obtained from one-step protein A purification.

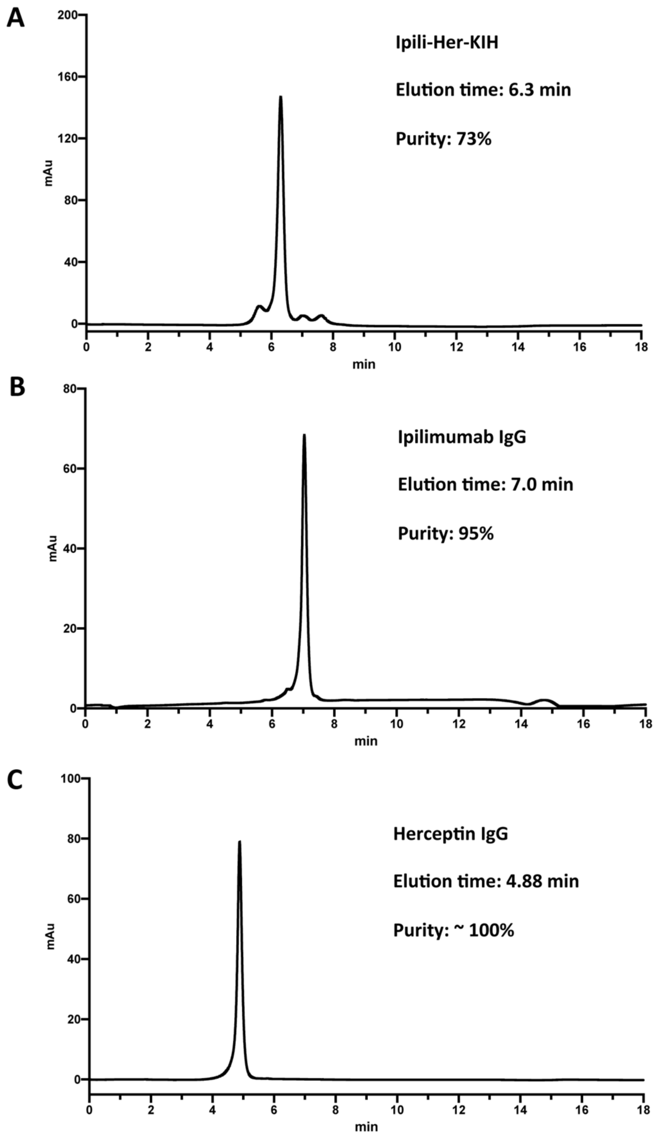


**Fig. S3 Purity assessment of the bispecific antibody Ipili-Her-KIH by analytical hydrophobic interaction chromatography.** The main peak of Ipili_Her_KIH (**A**) shows an elution time in-between of Ipilimumab (**B**) and Herceptin (**C**), representing the desired bispecific product that is estimated to be 73% of total protein obtained from one-step protein A purification.

**A**

| Estimated $EC_{50}$ for Fig. 2C, D | | |
|---|---|---|
| Antibody | $EC_{50}$ to CTLA4-Fc | $EC_{50}$ to CD38 |
| Ipilimumab | 1.10 ± 0.10 nM | |
| Ipili-Dara-KIH | 1.43 ± 0.11 nM | 1.44 ± 0.08 nM |
| Datatumumab | | 1.48 ± 0.18 nM |

**B**

| Estimated $EC_{50}$ for Fig. 2E, F | | |
|---|---|---|
| Antibody | $EC_{50}$ to CTLA4-Fc | $EC_{50}$ to ErbB2-Fc |
| Ipilimumab | 2.21 ± 0.13 nM | |
| Ipili-Her-KIH | 3.11 ± 0.24 nM | 0.72 ± 0.06 nM |
| Herceptin | | 1.58 ± 0.12 nM |

**Fig. S4 Assessment of ligand binding.** Estimated $EC_{50}$ values of bispecific antibodies, along with the original monoclonal antibodies, in ELISA-based binding study. $OD_{450}$ values were curve-fit (GraphPad Prism) to obtain $EC_{50}$ values.

**A**

| Apparent $K_d$ for antibody binding to antigen-expressing cells (Fig. 3E, F) | | |
|---|---|---|
| Antibody | $EC_{50}$ to Jurkat cells | $EC_{50}$ to MCF7 cells |
| Datatumumab | 0.31 ± 0.14 nM | |
| Ipilimumab | | weak |
| Ipili-Dara-KIH | 2.26 ± 0.26 nM | |
| Herceptin | | 0.60 ± 0.18 nM |
| Ipili-Her-KIH | | 1.39 ± 0.21 nM |

**B**

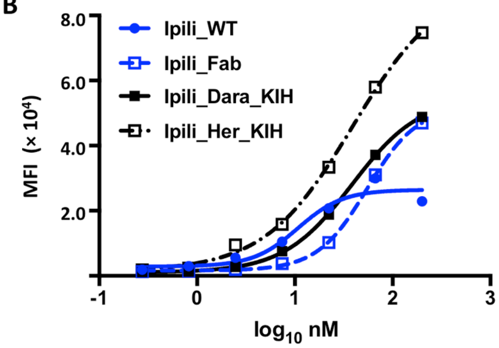


**C**

| Apparent $K_d$ for binding to CTLA4-expressing 293A cells | |
|---|---|
| Ipilimumab IgG | 9.23 ± 3.48 nM |
| Ipilimumab Fab | 101.30 ± 20.16 nM |
| Ipili-Dara-KIH | 47.10 ± 3.77 nM |
| Ipili-Her-KIH | 33.55 ± 3.06 nM |

**Fig. S5 Binding of bispecific antibodies to cell surface antigens.** (**A**) Binding to CD38 and ErbB2 was tested on Jurkat cells and MCF7 cells, respectively. The apparent $K_d$ values were obtained by curve fitting MFI values (Prism, GraphPad). (**B**) HEK293A cells were transfected with a plasmid expressing the human *CTLA4* gene, and binding by Ipili-Dara-KIH, Ipili-Her-KIH, or Ipilimumab IgG was assessed by flow cytometry. (**C**) The apparent $K_d$ values were obtained by curve fitting MFI values (Prism, GraphPad).

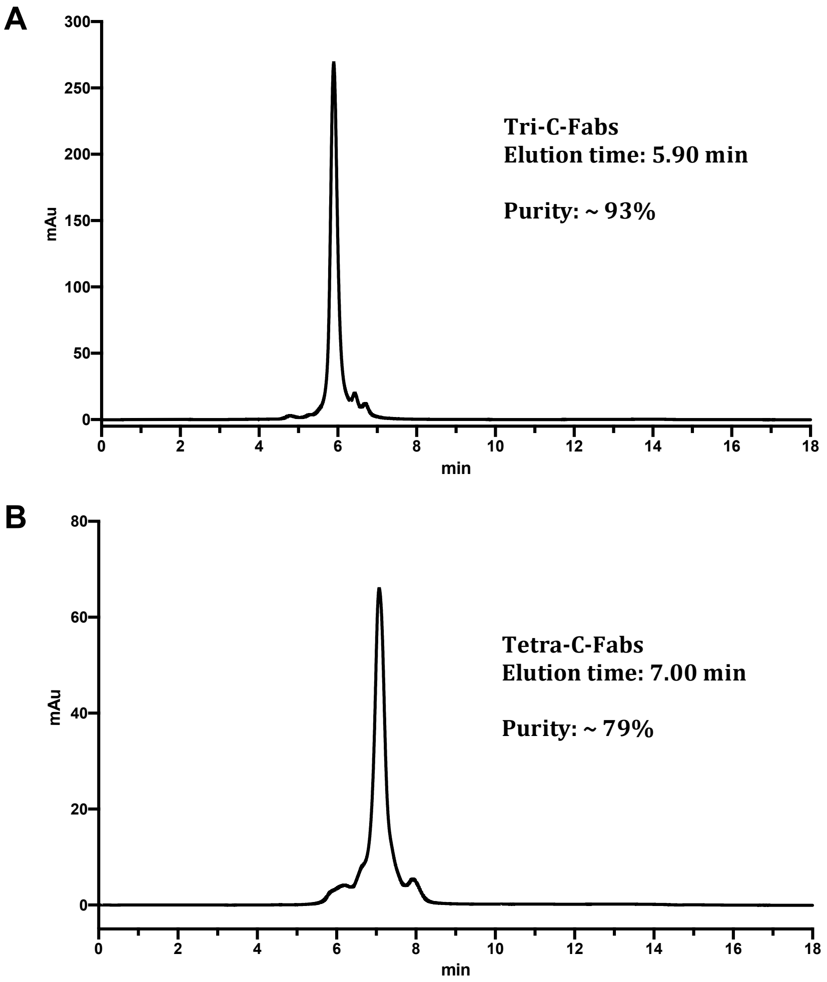


**Fig. S6 Purity assessment of tri-specific and tetra-specific antibodies by analytical hydrophobic interaction chromatography.** Purities of Tri-C-Fabs (**A**) and Tetra-C-Fabs (**B**) are estimated to be 93% and 79%, respectively.

**A**

| Estimated $EC_{50}$ for Fig.4 D, E, F | | | |
|---|---|---|---|
| Antibody | CTLA4-Fc | CD38 | ErbB2-Fc |
| Tri-N-Fabs | 0.85 ± 0.11 nM | 1.45 ± 0.13 nM | 0.67 ± 0.05 nM |
| Ipilimumab | 0.50 ± 0.06 nM | | |
| Daratumumab | | 1.49 ± 0.13 nM | |
| Herceptin | | | 0.27 ± 0.07 |

**B**

| Estimated $EC_{50}$ for Fig.5 D, E, F, G | | | | |
|---|---|---|---|---|
| Antibody | CTLA4-Fc | CD38 | ErbB2-Fc | PD-L1 |
| Tetra-N-Fabs | 0.86 ± 0.07 nM | 2.71 ± 0.26 nM | 0.25 ± 0.02 nM | 0.33 ± 0.03 nM |
| Ipilimumab | 0.49 ± 0.03 nM | | | |
| Daratumumab | | 1.47 ± 0.14 nM | | |
| Herceptin | | | 0.41 ± 0.03 | 0.40 ± 0.04 nM |

**C**

| Estimated $EC_{50}$ for Fig.6 D, E, F | | | |
|---|---|---|---|
| Antibody | CTLA4-Fc | CD38 | ErbB2-Fc |
| Tri-C-Fabs | 0.86 ± 0.11 nM | 1.36 ± 0.13 nM | 2.40 ± 0.26 nM |
| Ipilimumab | 0.50 ± 0.07 nM | | |
| Daratumumab | | 1.68 ± 0.17 nM | |
| Herceptin | | | 0.27 ± 0.03 |

**D**

| Estimated $EC_{50}$ for Fig.6 G, H, I, J | | | | |
|---|---|---|---|---|
| Antibody | CTLA4-Fc | CD38 | ErbB2-Fc | PD-L1 |
| Tetra-C-Fabs | 0.64 ± 0.07 nM | 1.73 ± 0.31 nM | 1.30 ± 0.08 nM | 0.45 ± 0.06 nM |
| Ipilimumab | 0.50 ± 0.04 nM | | | |
| Daratumumab | | 1.08 ± 0.17 nM | | |
| Herceptin | | | 0.36 ± 0.02 | 0.32 ± 0.03 nM |

**Fig. S7 Assessment of ligand binding of tri-specific and tetra-specific antibodies.** Estimated $EC_{50}$ values of ligand binding by Tri-N-Fabs (**A**), Tetra-N-Fabs (**B**), Tri-C-Fabs (**C**), Tetra-C-Fabs (**D**) along with the parental antibodies are analyzed by ELISA. $OD_{450}$ values were curve-fit (Prism, GraphPad) to obtain $EC_{50}$ values.

**A**

| Estimated $K_d$ values by biolayer-interferometry | | | | |
|---|---|---|---|---|
| | CTLA4-Fc | CD38 | ErbB2-Fc | PD-L1 |
| Ipilimumab | 3.17E-09 | | | |
| Ipilimumab Fab | 1.77E-08 | | | |
| Daratumumab | | 6.92E-10 | | |
| Herceptin | | | 9.95E-10 | |
| Atezolizumab | | | | 1.13E-09 |
| Ipili-Dara-KIH | 1.24E-08 | 5.59E-10 | | |
| Ipili-Her-KIH | 8.07E-09 | | 1.28E-09 | |
| Tri-N-Fabs | 1.06E-08 | 1.71E-08 | 3.26E-09 | |
| Tri-C-Fabs | 1.17E-08 | 1.40E-08 | 8.49E-09 | |
| Tetra-N-Fabs | 6.74E-09 | 7.95E-09 | 2.65E-09 | 3.80E-09 |
| Tetra-C-Fabs | 1.46E-08 | 2.76E-09 | 1.25E-08 | 7.14E-09 |

**B**

| Estimated $K_{off}$ values by biolayer-interferometry | | | | |
|---|---|---|---|---|
| | CTLA4-Fc | CD38 | ErbB2-FC | PD-L1 |
| Ipilimumab | 6.25E-04 | | | |
| Ipilimumab Fab | 1.77E-03 | | | |
| Daratumumab | | 1.03E-04 | | |
| Herceptin | | | 1.55E-04 | |
| Atezolizumab | | | | 2.75E-04 |
| Ipili-Dara-KIH | 1.58E-03 | 1.21E-04 | | |
| Ipili-Her-KIH | 1.20E-03 | | 1.98E-04 | |
| Tri-N-Fabs | 1.44E-03 | 1.25E-03 | 3.74E-04 | |
| Tri-C-Fabs | 2.04E-03 | 1.16E-03 | 6.81E-04 | |
| Tetra-N-Fabs | 1.53E-03 | 5.75E-04 | 6.87E-04 | 4.22E-04 |
| Tetra-C-Fabs | 1.64E-03 | 2.63E-04 | 6.81E-04 | 5.21E-04 |

**Fig. S8 Evaluation of antibody binding kinetics by biolayer-interferometry.** Estimated $K_d$ (**A**) and $K_{off}$ (**B**) values are summarized.
